# Analysis of Organophosphate Insecticide Half-Lives in Foods Fermented with Lactic Acid Bacteria

**DOI:** 10.64898/2026.02.04.703890

**Authors:** Julia Steenkamp, Graham Hepworth, Kate Howell

## Abstract

There is a growing concern that chronic, low-level exposure to organophosphate insecticides is a threat to human health. These synthetic chemicals are used in crop and livestock production all over the world, and the general population are exposed to them through consuming the residues that remain in food. Evidence is emerging that fermentation with lactic acid bacteria may be an effective way of reducing organophosphate insecticide residues. However, while several studies have investigated this topic, outcome measures have varied, and there has been no research to date which has consolidated this data to better understand the half-lives of organophosphate insecticides in fermented foods and the factors affecting degradation. The aim of this review was to synthesise the evidence on organophosphate insecticide degradation during lactic acid fermentation, and analyse organophosphate insecticide half-lives, in order to determine the effectiveness of lactic acid fermentation in reducing organophosphate insecticide residues in food. Furthermore, the study aimed to explore the factors that impact the rate of degradation. CAB Abstracts, Food Science and Technology Abstracts, Scopus and Web of Science were searched for eligible laboratory-based studies, which were published after 2000. The literature search and screening process resulted in the inclusion of 14 eligible studies. Studies were screened for Risk of Bias (ROB) using the RoBDMAT tool. Collated results showed that organophosphate insecticides degraded over time, and this was irrespective of fermentation. However, out of the 249 experiments that involved a controlled fermentation, 232 demonstrated that fermentation with lactic acid bacteria could speed up the degradation of organophosphate insecticides in food, beyond the rate of inherent degradation in the food matrix, leading to shorter half-lives. The half-lives of organophosphate insecticides in apple juice, milk and wheat ranged from 9.5 hours to 21 days in fermented foods and ranged from 21.4 hours to 36.5 days in non-fermented foods. Single species of lactic acid bacteria that demonstrated strong potential for organophosphate insecticide degradation were *Lpb.plantarum* subsp*.plantarum, Lab.delbrueckii* subsp*.bulgaricus* and *Lvb. brevis,* where the median percentage change in organophosphate insecticide half-life during fermentation was -42.3%, -25.0% and -22.9%, respectively. Organophosphate insecticide degradation during natural fermentation was less clear because of fewer studies and less consistent results. Whilst the collated data shows that fermentation with lactic acid bacteria is an effective method to reduce organophosphate insecticide residues in food, reflected in shorter half-lives, the small number of studies and variability among studies does limit the conclusions that can be drawn, and further research is needed to strengthen these findings. The results of our analysis may help to inform more reliable organophosphate exposure assessments for the population as well as provide novel insights for both consumers and food manufacturers, expanding the market potential for fermented foods.

## Introduction

In the 1940s the world of agriculture was forever changed by the introduction of synthetic pesticides. Discovery of chemicals such as Dichlorodiphenyltrichloroethane (DDT) brought with them the promise of effective insect control in crop and livestock production. Alongside the introduction of high-yield crop varieties, synthetic pesticides were soon recognised as a major part of the solution to feeding a growing population. However, by the 1970s it was clear that these organochlorine pesticides were a major threat to the environment and human health. And so, alternative synthetic pesticides, such as organophosphates, which were considered safer and less environmentally persistent, slowly replaced organochlorines (Tudi et al., 2021).

Large scale synthesis of organophosphates began during World War II where they were used as both chemical warfare agents and a means to protect soldiers from disease (Boczkowski et al., 2025; De Castro et al., 2017). They soon became one of the most widely used pesticides globally and continue to be used today on a wide variety of crops (Australian Pesticides and Veterinary Medicines Authority, 2025; Fagundes et al., 2025).

Organophosphate insecticides are esters of phosphoric, phosphonic, phosphinic or phosphoramidic acids; designed to kill insects through the inhibition of acetylcholine esterase (Jaiswal et al., 2024). When organophosphate insecticides are applied to food crops, only a small portion reaches the target organism, with the remainder persisting in the soil and on the surface of the crop, and also dispersing into the surrounding environment (Kiruthika et al., 2025; Perpetuini et al., 2023; Sarlak et al., 2021). The primary route of exposure to organophosphate insecticides for the general population is through ingestion of residues in food (Chen et al., 2023; Jaiswal et al., 2024; Perpetuini et al., 2023).

Organophosphate insecticide residues have been detected in the Australian food system (Food Standards Australia New Zealand, 2019), and there is evidence these insecticides make their way into the human body, with studies demonstrating that their metabolites have been found in the urine samples of Australian adults and children (Babina et al., 2012; English et al., 2019; Heffernan et al., 2016; Y. Li et al., 2019). Organophosphate insecticides are a threat to human health. Research on the long-term effects of chronic exposure to these insecticides is emerging, with the impact on childhood neurodevelopment of particular concern (Farkhondeh et al., 2020).

Whilst organophosphate insecticides are known to slowly degrade over time, various food processing methods are able to either remove the insecticides or speed up their degradation; including peeling, washing and thermal processing as well as the use of chemicals such as chlorine dioxide and ozone, and technologies such as pulsed electric field, irradiation, hydrostatic pressure and ultra sonification (Mir et al., 2022). There is a growing interest in the role of microbes in the degradation of organophosphate insecticides. Most research in this area has been in soil bioremediation, rather than in food. The first known organophosphate insecticide degrading bacteria was isolated from the rice fields of the Philippines and was identified as *Flavobacterium sp*. Since that time, several organophosphate degrading bacteria have been isolated and identified from soils around the world (Mali et al., 2023). More recently, researchers have begun investigating whether bacteria can perform this same function in foods contaminated with organophosphate insecticides.

Lactic acid bacteria are a diverse group of gram-positive, non-spore forming, aerotolerant microorganisms, predominantly of the order Lactobacillales (Kiruthika et al., 2025; Rossi, 2023; Wang et al., 2021; Zapaśnik et al., 2022). These bacteria have the ability to transform the taste, appearance, texture, preservation properties and nutritional qualities of food through fermentation, and are responsible for producing some of the commonly consumed fermented foods from around the world, such as gari in Nigeria, kimchi in Korea, olives in Spain, kefir in Russia, sauerkraut in Germany and yoghurt in Australia (Cuamatzin-García et al., 2022; Wang et al., 2021).

Evidence is emerging that during the fermentation process, lactic acid bacteria may also contribute to the degradation of organophosphate insecticides in food (Kiruthika et al., 2025). Researchers have studied a variety of different foods contaminated with an array of organophosphate insecticides. Furthermore, numerous species of lactic acid bacteria have been investigated under various incubation conditions. Outcome measures have varied between studies and there has been no research to date which has consolidated the evidence to better understand organophosphate insecticide half-lives in fermented foods and the factors affecting degradation. Therefore, the purpose of this review was to synthesise the evidence on organophosphate insecticide degradation during lactic acid fermentation, and analyse organophosphate insecticide half-lives, in order to determine the effectiveness of lactic acid fermentation in reducing organophosphate insecticide residues in food. Additionally, we sought to explore the factors that impact the rate of degradation.

A search strategy was developed, and the literature was systematically searched for studies that met the selection criteria. Studies were included that involved food being spiked with an organophosphate insecticide in the laboratory and then subjected to a lactic acid fermentation (controlled or natural), with insecticide residues measured before, during and/or immediately after the incubation period. An exploratory look at the data with descriptive statistics was considered more appropriate than a meta-analysis due to substantial methodological heterogeneity in studies relating to organophosphate insecticide type and contaminated level, lactic acid bacteria species and inoculation concentration, as well as fermentation conditions. This heterogeneity does limit how well the results represent a true effect, and subsequently the conclusions that can be drawn. Nonetheless, the analysis of these studies demonstrated that, in most cases, controlled fermentation with lactic acid bacteria, shortened the half-life of organophosphate insecticides in food.

This knowledge will help to inform consumers of the types of foods and food processes that can lower insecticide exposure. Having a better understanding of the role of fermentation in reducing organophosphate insecticides may also increase the market potential for fermented foods and assist food manufactures in tailoring fermentation processes for optimum organophosphate insecticide degradation. This knowledge may also lead to the development of more accurate food processing factors for fermentation, to inform pesticide exposure assessments for the protection of public health.

## Methods

### Eligibility Criteria

Studies were included in the review based on the following inclusion criteria:

### Population

The study was carried out in food contaminated with organophosphate insecticides. These insecticides had either been applied during production or spiked in the laboratory.

### Intervention

The intervention was a fermentation process involving lactic acid bacteria. Lactic acid bacteria are microorganisms of the order Lactobacillales, which is comprised of six families. It is noted that in 2020 a new taxonomy was released for the Lactobacillaceae family (Zheng et al., 2020). Lactobacillus was re-classified into 25 genera, and an additional 23 novel genera were added. The new taxonomy was applied to all species included in this review. A fermentation involving lactic acid bacteria was evident through:

▪ Inoculation of food with lactic acid bacteria with conditions conducive to fermentation, or
▪ Inoculation of milk with a commercial yoghurt starter culture with conditions conducive to fermentation, or
▪ For natural fermentations, isolation and identification of lactic acid bacteria and measurement of pH, with conditions conducive to fermentation.

### Comparator

Comparators were negative controls which had not undergone a fermentation but were otherwise exposed to the same conditions. For controlled fermentations, controls were not inoculated with lactic acid bacteria or commercial yoghurt starter culture. For natural fermentations, controls were not exposed to conditions conducive to fermentation.

### Outcome Measures

Studies were included which reported one or more of the following:

▪ Organophosphate insecticide concentration before and after fermentation
▪ Percentage degradation of organophosphate insecticide following fermentation
▪ Organophosphate degradation rate constant (k)
▪ Organophosphate insecticide half-life (T_1/2_)

Exploratory data was also collected on the pH of treatment and control samples during fermentation as well as the type and concentration of organophosphate insecticide degradation products that formed during fermentation.

### Study Type

Included studies were laboratory experiments.

### Information Sources & Search Strategy

Four databases were searched, including CAB Abstracts, Food Science and Technology Abstracts (FSTA), Scopus and Web of Science (WOS). The search strategy was built around three concepts: Organophosphate insecticides, lactic acid fermented foods, and the lactic acid fermentation process. Based on this, the following search string was developed:

(organophosph* OR acephate OR azamethiphos OR azinphos OR bensulide OR bromophos OR cadusafos OR chlorfenvinphos OR chlorpyrifos OR chlorthiophos OR coumaphos OR crotoxyphos OR demeton OR dialifos OR diazinon OR dichlofenthion OR dichlorvos OR dicrotophos OR dimethoate OR dioxathion OR disulfoton OR ethephon OR ethion OR ethoprophos OR famphur OR fenamiphos OR fenchlorphos OR fenitrothion OR fensulfothion OR fenthion OR formothion OR isofenphos OR leptophos OR malathion OR menazon OR methacrifos OR methamidophos OR methidathion OR mevinphos OR monocrotophos OR naled OR naphthalophos OR omethoate OR oxydemeton OR parathion OR phenthoate OR phorate OR phosalone OR phosfolan OR phosmet OR phoxim OR pirimiphos OR profenofos OR prothiofos OR pyrazophos OR sulprofos OR temephos OR terbufos OR tetrachlorvinphos OR thiometon OR trichlorfon) AND (food OR milk OR yoghurt OR yogurt OR cheese OR kefir OR lassi OR cabbage OR sauerkraut OR kraut OR kimchi OR pickl OR wheat OR sourdough OR meat OR salami OR pepperoni OR chorizo OR ferment*) AND (ferment* OR “lactic acid bacteria” OR lactobac* OR yoghurt OR yogurt OR cheese OR kefir OR lassi OR sauerkraut OR kraut OR kimchi OR pickl OR sourdough OR salami OR pepperoni OR chorizo).

The search was refined by eliminating citations where the following words or phrases were present in the title: soil OR silage OR feed OR flame OR plastic* OR “chemical weapon” OR “chemical warfare” OR “nerve agent” OR toxic* OR poison* OR animal* OR human* OR adult*.

The search was limited to papers published from 2000 onwards.

### Selection Process

One reviewer collected and screened each citation. Where citations did not clearly meet the inclusion criteria, a second reviewer was consulted.

### Quality Appraisal

Included studies were assessed for Risk of Bias (RoB) using the RoBDMAT tool (Delgado et al., 2022). This tool was selected as RoB tools for use in laboratory studies in food science and microbiology are not available; and the RoBDMAT tool, despite being developed for studies in dentistry, is suitable for use in other non-human, non-animal studies.

The tool assesses risk of bias across four domains: 1) Planning and allocation, 2) Sample/specimen preparation, 3) Outcome assessment, and 4) Data treatment and outcome reporting. Nine signalling questions pertaining to bias within each domain can be answered with either sufficiently reported / adequate, insufficiently reported, not reported / not adequate or not applicable. The tool does not generate a ‘final score’ as this would involve attributing weights to different domains which is subjective and not recommended.

In addition to the RoB assessment, further elements of the study methodology were evaluated to determine robustness and alignment with the objective of the review. These elements related to:

▪ The measurement of pH in treatment and control samples throughout the fermentation period
▪ The analysis of organophosphate degradation products during fermentation
▪ Whether a standard fermentation procedure was followed, in line with the Australian Food Standards Code (Food Standards Australia New Zealand, 2025) and the Fermented Foods Safety Guidance (Hudson et al., 2024).

### Data Items

Organophosphate insecticide half-lives were recorded for insecticides in food during fermentation (treatment samples) and for insecticides in food without fermentation (control samples). If this data was not available, it was calculated from degradation rate constants (K), which were either directly reported or calculated from experimental data.

#### Calculating Degradation Rate Constants (K) from experimental data

A first-order degradation model was used to calculate degradation rate constants. This was based on the assumption that organophosphate insecticide degradation satisfies pseudo first-order degradation kinetics, whereby a constant proportion of insecticide is degraded per unit of time (Cardozo et al., 2019). The first order reaction model is expressed below:

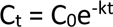

Where C_t_ is the concentration of organophosphate insecticide (mg/kg) at time t (hours or days), C_0_ is the initial concentration of organophosphate insecticide (mg/kg) and k is the degradation rate constant (/hour or /day).

Degradation rate constants were determined from experimental data by plotting ln(C_t_) versus time and determining the slope of the line (k).

#### Calculating Pesticide Half-Lives from Degradation Rate Constants (K)

Organophosphate insecticide half-lives were calculated using the first order reaction model by substituting Ct= ½ C_0_. The formula for half-life is expressed below:

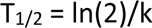

Where T_1/2_ is the half-life of the organophosphate insecticide in hours or days.

When organophosphate insecticide half-lives were calculated, the Coefficient of Determination, R-squared (R^2^), was analysed to determine how well the data conformed to the first-order degradation model. For experiments where the data did not fit well (reflected in a low R^2^) organophosphate insecticide degradation was instead expressed as a percentage degradation over the fermentation period, reflecting the amount of insecticide degraded rather than a degradation rate.

#### Effect Measures

The half-lives of organophosphate insecticides in fermented food samples (treatment) and non-fermented food samples (control) were compared. The difference was expressed as a percentage change, using the following formula:

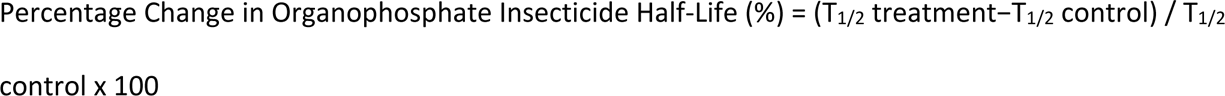

Where T_1/2_ is the half-life of the organophosphate insecticide in hours or days. Data synthesis

A narrative synthesis was presented whereby results were categorised according to fermentation type. For the purpose of this review, ‘controlled fermentation’ was used to describe a fermentation process involving initial heat treatment followed by deliberate inoculation with lactic acid bacteria and subsequent incubation. ‘Natural fermentation without inoculation’ was used to describe a fermentation process occurring without inoculation with lactic acid bacteria, relying solely on the activity of lactic acid bacteria naturally present within the food. ‘Natural fermentation with inoculation’ was used to describe a fermentation process relying on the activity of lactic acid bacteria naturally present within food, as well as an additional inoculation of lactic acid bacteria. A meta-analysis was not conducted due to the substantial methodological heterogeneity of the data in relation to organophosphate insecticide type, organophosphate insecticide contamination level, lactic acid bacteria species, inoculation concentration and fermentation conditions.

### Statistics

Data was analysed using SPSS version 3.0. Results were expressed showing means and 95% confidence intervals when comparing half-lives, and medians and interquartile ranges when comparing percentage change in half-lives. Kinetic parameters were calculated with linear regression analysis.

## Results

### Study Selection

The search yielded 987 citations. After removal of duplicates, 698 citations were sorted for retrieval. Following screening of titles, abstracts and full-text, 14 studies met the inclusion criteria and were included in the review. The PRISMA flow diagram, outlining the results of the screening process, is represented in Figure 1.

**Figure 1.**
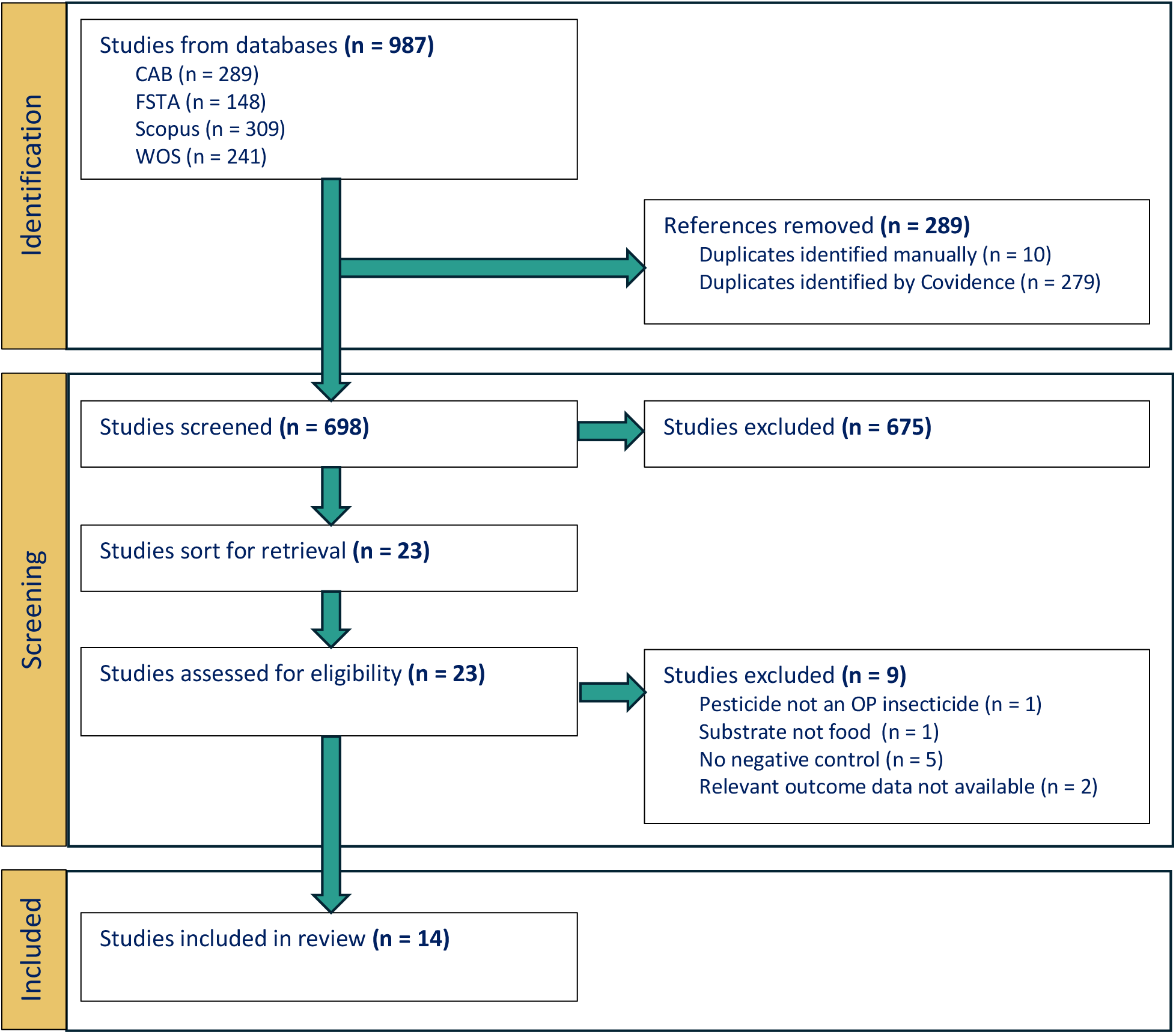
PRISMA Flow Diagram outlining the results of the screening process. CAB = CAB Abstracts, FSTA = Food Science and Technology Abstracts, WOS = Web of Science. n = number of studies.

### Characteristics of Included Studies

The data from 14 studies encompassing 277 fermentations was synthesised, of which 249 were controlled fermentations (Figure S1). The characteristics of these studies are represented in Table 1. Studies were undertaken in China, Iran, Serbia and Turkey between 2011 and 2024. Whilst studies in milk were most common, other fermentation substrates included apple juice, cabbage, olives and wheat. Fourteen distinct organophosphate insecticides were investigated, with organophosphate insecticide contamination levels between 0.5mg/kg and 50mg/kg. The most studied organophosphate insecticides were chlorpyrifos-methyl, diazinon and malathion (Figure S1). All insecticides were applied to food in the laboratory, post-harvest, rather than during production. Ten species of lactic acid bacteria were investigated, with fermentation temperatures ranging between 5°C and 45^°^C and fermentation times ranging from 5 hours to 60 days. The most studied lactic acid bacteria were *Lpb. plantarum* subsp*.plantarum* and *Lab. delbrueckii* subsp*.bulgaricus* (Figure S1).

**Table 1.**
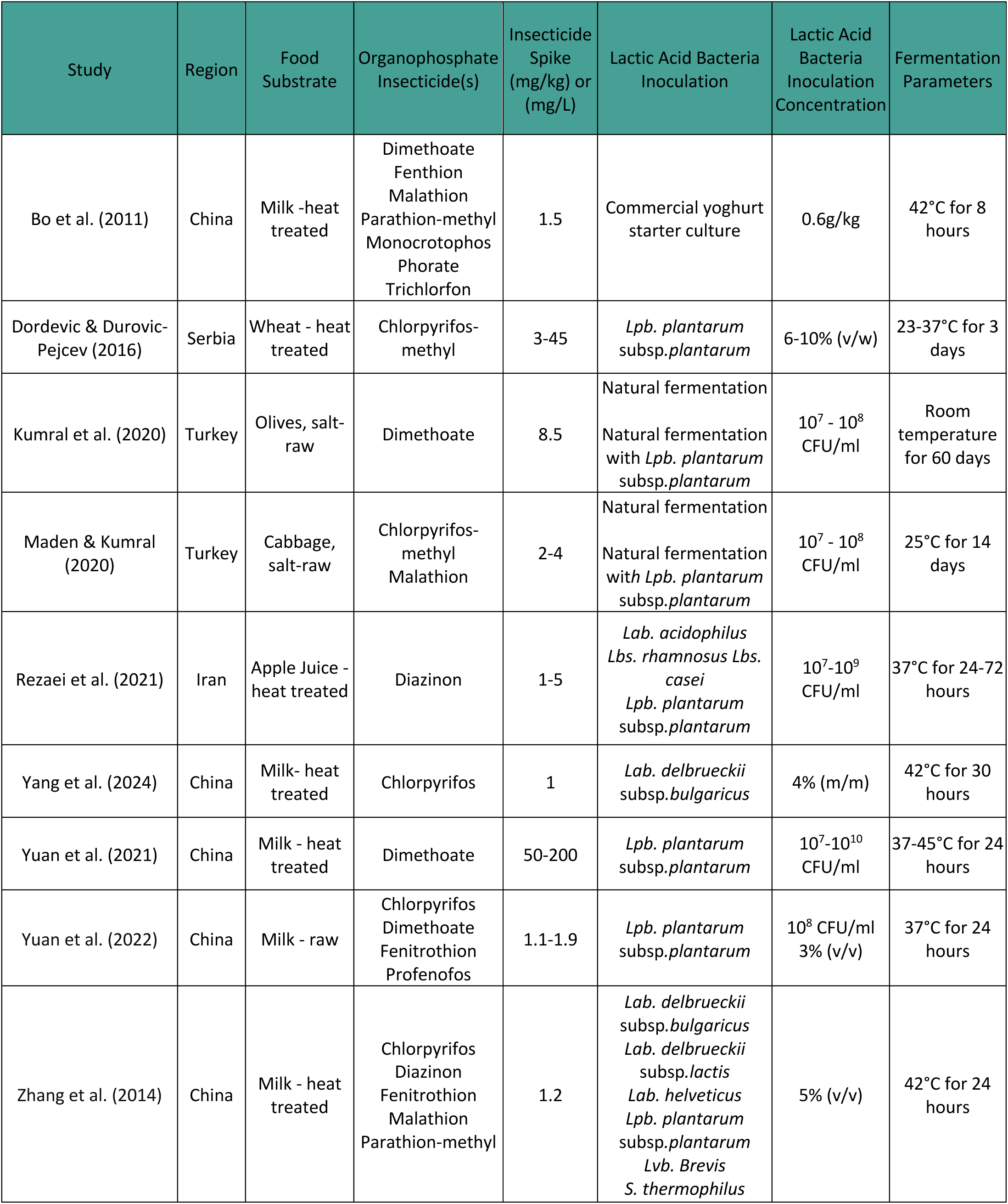
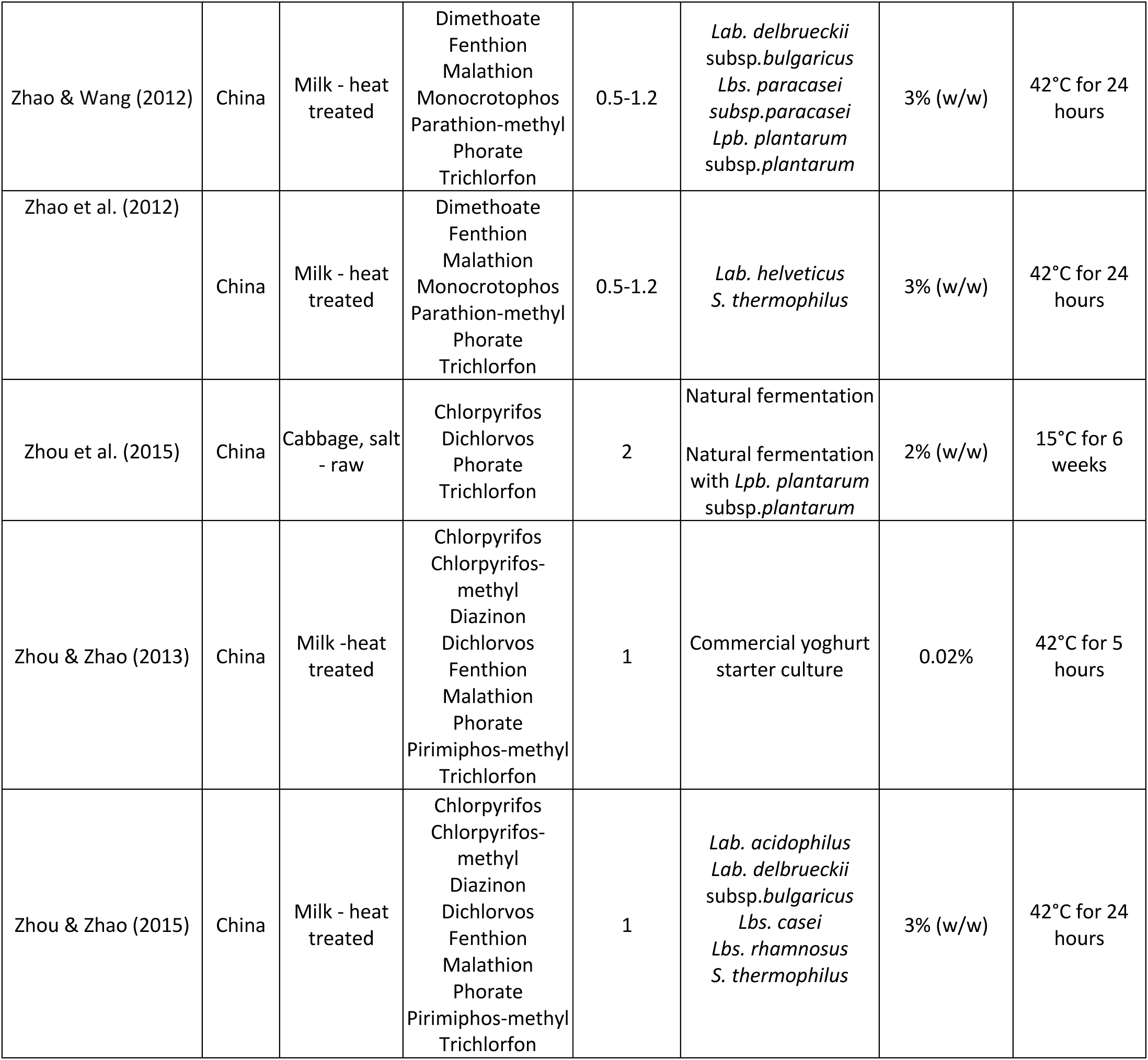
Characteristics of included studies. The characteristics of the 14 included studies are outlined. ‘Heat treated’ indicates that the food was pasteurised or sterilised prior to fermentation. ‘Insecticide spike’ refers to the concentration of organophosphate insecticide applied to food in the laboratory prior to fermentation. Commercial yoghurt starter culture typically contained a combination of *Lab. delbrueckii* subsp.b*ulgaricus* and *S. thermophilus* but this was not always specified. ‘Fermentation parameters’ refers to the temperature and length of the fermentation incubation period. v/w = volume of lactic acid bacteria per weight of food, expressed as a percentage. CFU/ml = colony forming units per millilitre.

| Study | Region | Food Substrate | Organophosphate Insecticide(s) | Insecticide Spike (mg/kg) or (mg/L) | Lactic Acid Bacteria Inoculation | Lactic Acid Bacteria Inoculation Concentration | Fermentation Parameters |
| --- | --- | --- | --- | --- | --- | --- | --- |
| Bo et al. (2011) | China | Milk -heat treated | Dimethoate<br>Fenthion<br>Malathion<br>Parathion-methyl<br>Monocrotophos<br>Phorate<br>Trichlorfon | 1.5 | Commercial yoghurt starter culture | 0.6g/kg | 42°C for 8 hours |
| Dordevic & Durovic-Pejcev (2016) | Serbia | Wheat - heat treated | Chlorpyrifos-methyl | 3-45 | *Lpb. plantarum* subsp.*plantarum* | 6-10% (v/w) | 23-37°C for 3 days |
| Kumral et al. (2020) | Turkey | Olives, salt-raw | Dimethoate | 8.5 | Natural fermentation<br>Natural fermentation with *Lpb. plantarum* subsp.*plantarum* | 10<sup>7</sup> - 10<sup>8</sup> CFU/ml | Room temperature for 60 days |
| Maden & Kumral (2020) | Turkey | Cabbage, salt-raw | Chlorpyrifos-methyl<br>Malathion | 2-4 | Natural fermentation<br>Natural fermentation with *Lpb. plantarum* subsp.*plantarum* | 10<sup>7</sup> - 10<sup>8</sup> CFU/ml | 25°C for 14 days |
| Rezaei et al. (2021) | Iran | Apple Juice - heat treated | Diazinon | 1-5 | *Lab. acidophilus*<br>*Lbs. rhamnosus* *Lbs. casei*<br>*Lpb. plantarum* subsp.*plantarum* | 10<sup>7</sup>-10<sup>9</sup> CFU/ml | 37°C for 24-72 hours |
| Yang et al. (2024) | China | Milk- heat treated | Chlorpyrifos | 1 | *Lab. delbrueckii* subsp.*bulgaricus* | 4% (m/m) | 42°C for 30 hours |
| Yuan et al. (2021) | China | Milk - heat treated | Dimethoate | 50-200 | *Lpb. plantarum* subsp.*plantarum* | 10<sup>7</sup>-10<sup>10</sup> CFU/ml | 37-45°C for 24 hours |
| Yuan et al. (2022) | China | Milk - raw | Chlorpyrifos<br>Dimethoate<br>Fenitrothion<br>Profenofos | 1.1-1.9 | *Lpb. plantarum* subsp.*plantarum* | 10<sup>8</sup> CFU/ml<br>3% (v/v) | 37°C for 24 hours |
| Zhang et al. (2014) | China | Milk - heat treated | Chlorpyrifos<br>Diazinon<br>Fenitrothion<br>Malathion<br>Parathion-methyl | 1.2 | *Lab. delbrueckii* subsp.*bulgaricus*<br>*Lab. delbrueckii* subsp.*lactis*<br>*Lab. helveticus*<br>*Lpb. plantarum* subsp.*plantarum*<br>*Lvb. Brevis*<br>*S. thermophilus* | 5% (v/v) | 42°C for 24 hours |
| Zhao & Wang (2012) | China | Milk - heat treated | Dimethoate<br>Fenthion<br>Malathion<br>Monocrotophos<br>Parathion-methyl<br>Phorate<br>Trichlorfon | 0.5-1.2 | *Lab. delbrueckii* subsp.*bulgaricus*<br>*Lbs. paracasei* subsp.*paracasei*<br>*Lpb. plantarum* subsp.*plantarum* | 3% (w/w) | 42°C for 24 hours |
| Zhao et al. (2012) | China | Milk - heat treated | Dimethoate<br>Fenthion<br>Malathion<br>Monocrotophos<br>Parathion-methyl<br>Phorate<br>Trichlorfon | 0.5-1.2 | *Lab. helveticus*<br>*S. thermophilus* | 3% (w/w) | 42°C for 24 hours |
| Zhou et al. (2015) | China | Cabbage, salt - raw | Chlorpyrifos<br>Dichlorvos<br>Phorate<br>Trichlorfon | 2 | Natural fermentation<br>Natural fermentation with *Lpb. plantarum* subsp.*plantarum* | 2% (w/w) | 15°C for 6 weeks |
| Zhou & Zhao (2013) | China | Milk -heat treated | Chlorpyrifos<br>Chlorpyrifos-methyl<br>Diazinon<br>Dichlorvos<br>Fenthion<br>Malathion<br>Phorate<br>Pirimiphos-methyl<br>Trichlorfon | 1 | Commercial yoghurt starter culture | 0.02% | 42°C for 5 hours |
| Zhou & Zhao (2015) | China | Milk - heat treated | Chlorpyrifos<br>Chlorpyrifos-methyl<br>Diazinon<br>Dichlorvos<br>Fenthion<br>Malathion<br>Phorate<br>Pirimiphos-methyl<br>Trichlorfon | 1 | *Lab. acidophilus*<br>*Lab. delbrueckii* subsp.*bulgaricus*<br>*Lbs. casei*<br>*Lbs. rhamnosus*<br>*S. thermophilus* | 3% (w/w) | 42°C for 24 hours |

### Quality Appraisal

All 14 studies were assessed for risk of bias using the RoBDMAT tool and the results are represented in Figure 2. As per the selection criteria, all studies included a negative control. However, Yuan and colleagues (2021) did not include a negative control for all organophosphate insecticide spike levels. Only those experiments that included a negative control were included in the analysis of controlled fermentations. In relation to randomisation, nine studies used a randomisation process whereby a small selection of samples were randomly selected from bulk samples for analysis. In each of these studies, the full details of the randomisation process were not reported. Remaining studies either prioritised standardisation over randomisation or did not randomise despite there being likely variability between samples. All 14 studies did not report a sample size rationale, with the majority of studies performing all experiments in triplicate. Blinding of the test operator was not reported in any of the 14 studies. Eight studies were assessed as ‘insufficiently reported’ for statistical analysis. This was either because statistical methods were not adequately reported or because there were no statistical tests performed to substantiate differences between groups. This may have been a reflection of differences in study objectives and hypotheses.

**Figure 2.**
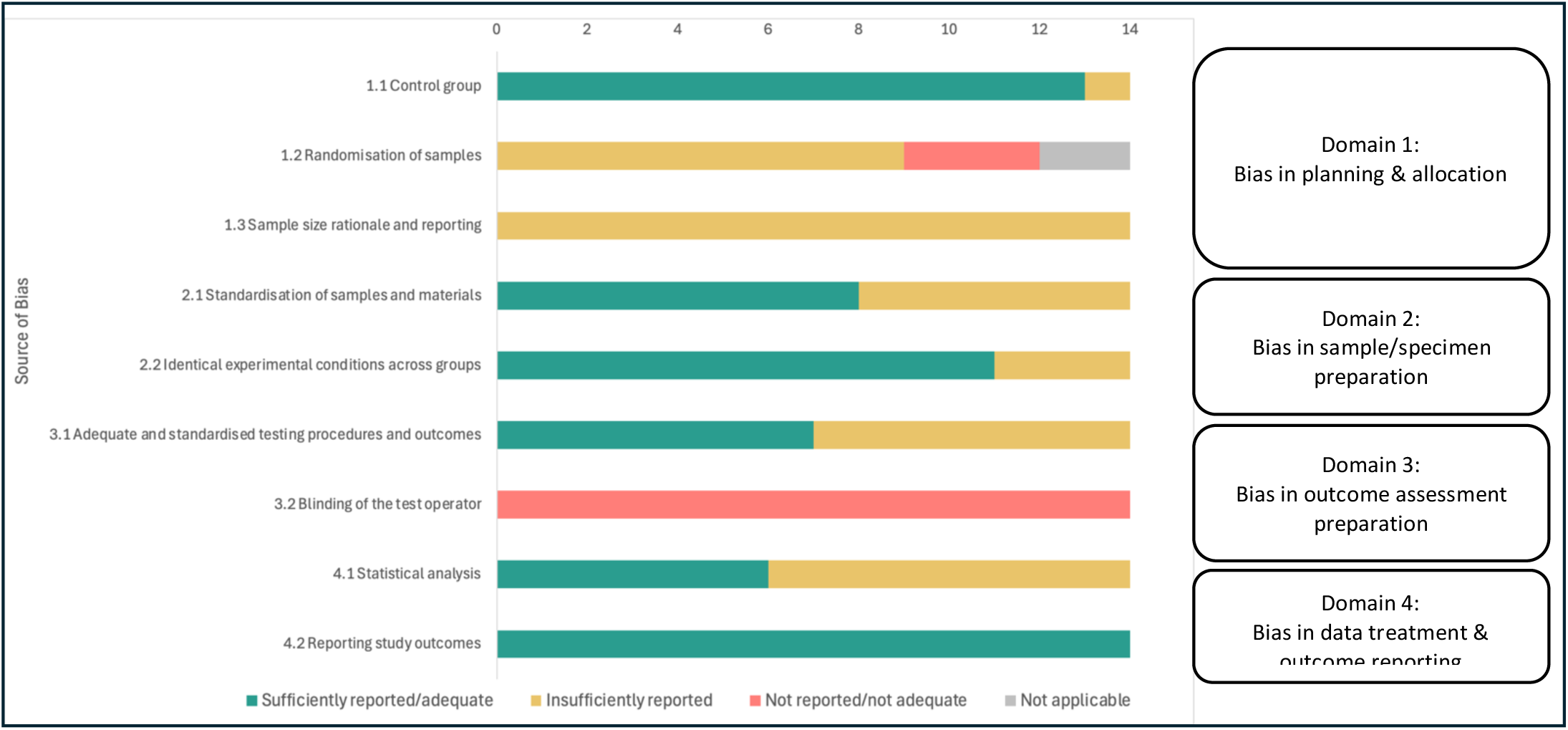
Results of RoBDMAT risk of bias assessment for 14 included studies. 14 studies underweight a risk of bias assessment using the RoBDMAT tool. The tool includes four domains: 1) Bias in planning and allocation, 2) Bias in sample / specimen preparation, 3) Bias in outcome assessment, and 4) Bias in data treatment and outcome reporting. Nine signalling questions pertaining to bias within each domain were answered with sufficiently reported / adequate (green), insufficiently reported (amber), not reported / not adequate and not applicable (red). ‘Not applicable’ (grey) was used for ‘1.2 randomisation of samples’ when a study prioritised standardisation of samples over randomisation and the nature of the samples did not imply variability.

Further evaluation of study designs identified that six studies measured pH, but only two of these measured pH in both treatment and control samples (Maden & Kumral, 2020; Rezaei et al., 2021). Two studies measured the concentration of organophosphate degradation products throughout the fermentation period (Yang et al., 2024; Yuan et al., 2021). Thirteen studies followed a standard fermentation procedure. One study in milk did not follow a standard fermentation procedure, as a heat treatment was not carried out prior to inoculation with lactic acid bacteria (Yuan et al., 2022). Whilst this was appropriate for the objectives of the study, the study was not included in this analysis as it did not fit the definition of ‘controlled fermentation’ and the results would likely vary for heat-treated versus non heat-treated milk (Bamforth & Cook, 2019).

### Organophosphate Insecticides Degrade over Time

Across 13 studies, it was observed that organophosphate insecticides degraded over time, and this was irrespective of fermentation. The half-lives of organophosphate insecticides in apple juice, milk and wheat, were all less than 37 days, demonstrating that these insecticides degraded during the incubation period, regardless of whether they were inoculated with lactic acid bacteria. This was also observed in olives and cabbage, where organophosphate insecticides degraded by at least 10% over the incubation period in treatment and control samples. Yuan and colleagues (2022) observed an exception to this trend in their study involving unpasteurised, raw milk. Whilst fenitrothion, chlorpyrifos, profenfos and dimethoate degraded by up to 79% in raw milk which had been inoculated with *Lpb. plantarum* subsp*.plantarum*, all four insecticides showed negligible degradation in uninoculated samples, persisting in the raw milk through the incubation period.

### Controlled Fermentation Shortens the Half-Life of Organophosphate Insecticides

Ten studies involving 249 treatments investigated organophosphate insecticide degradation during controlled fermentation, and these were in contaminated apple juice, milk and wheat (Figure S1). In these studies, degradation fitted well with the first-order degradation model and half-lives were compared between treatment and control samples.

The half-lives of organophosphate insecticides, on average, were shorter in fermented foods compared with non-fermented foods (60.2 hours, SD = 66.0 vs. 135.1 hours, SD = 172.7 hours, respectively), and this trend was observed across all three foods (Figure 3). It is noted that wide confidence intervals indicated a level of uncertainty in these results, a reflection of variability in the data and, in some cases, a small number of experiments.

**Figure 3.**
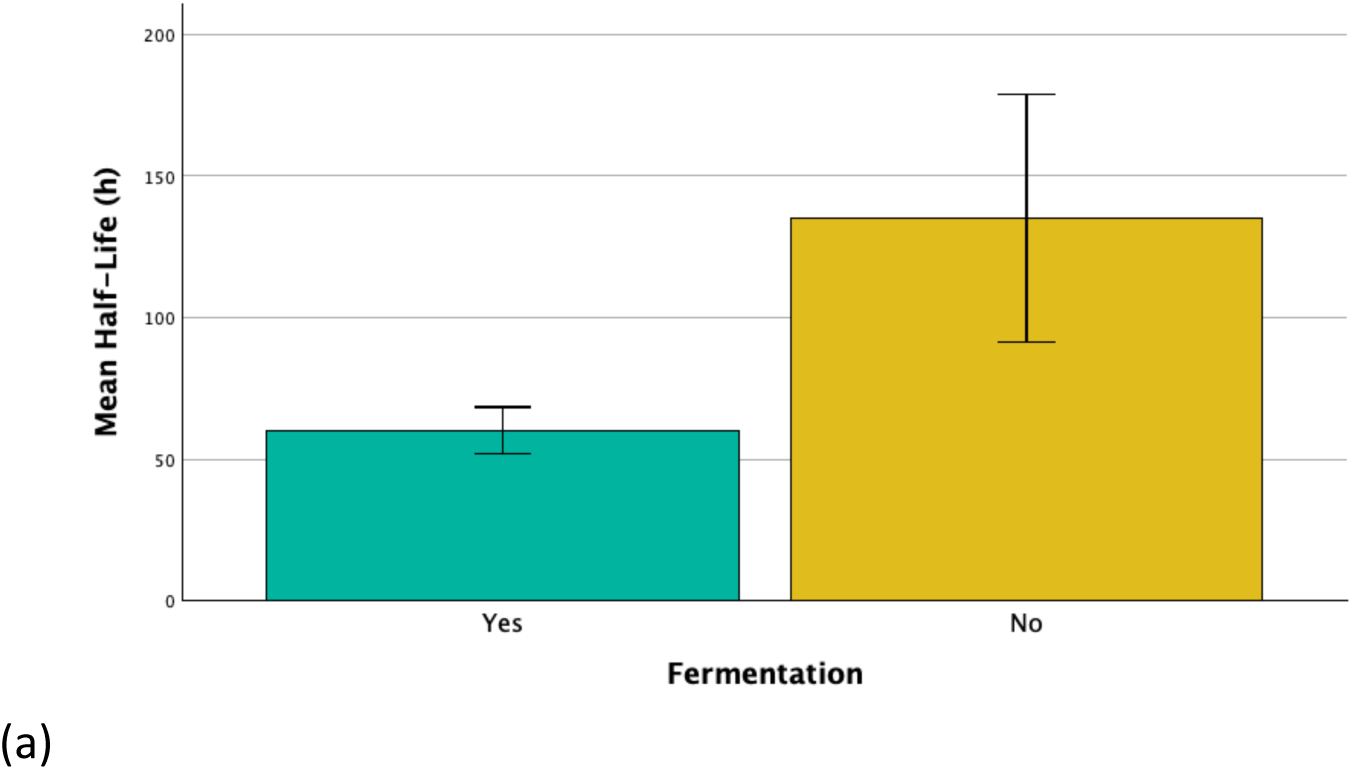
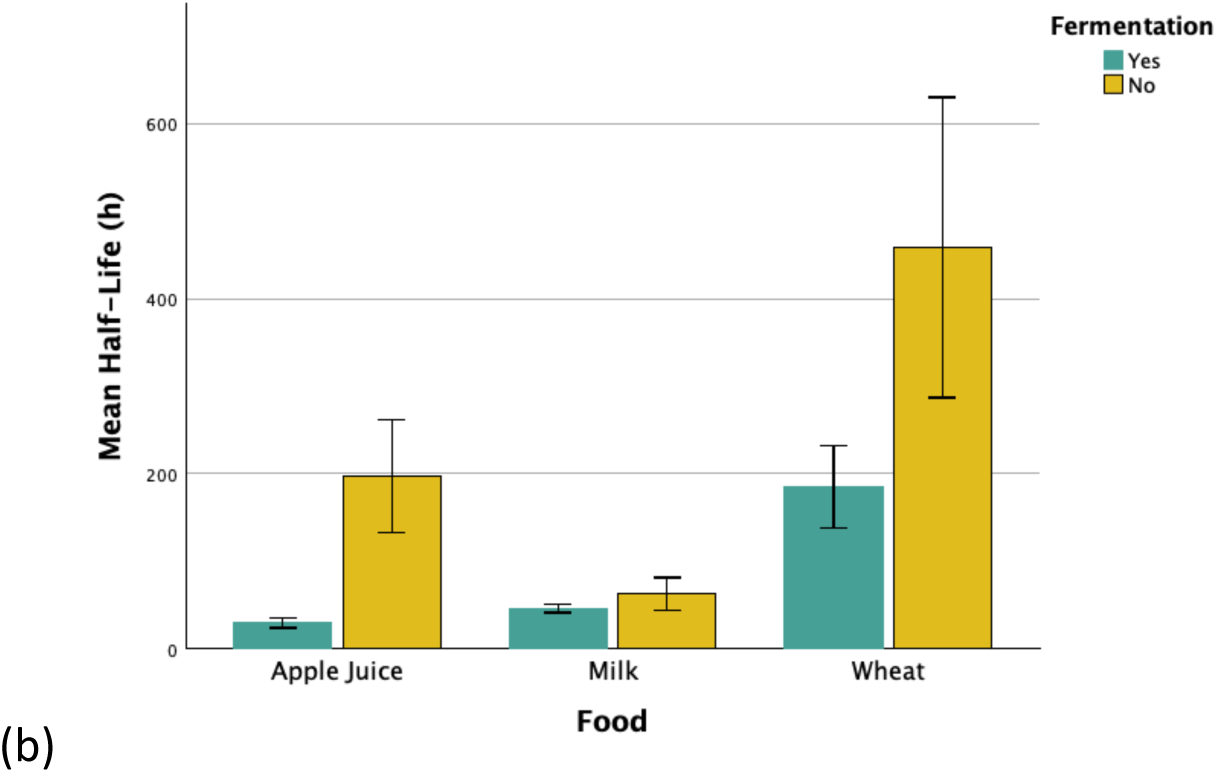
Mean organophosphate insecticide half-life during controlled fermentation compared with no fermentation. Combined results of 10 studies shown in (a) and for each of the food groups separately in (b). ‘Fermentation = Yes’ represents treatment groups (fermented foods), whereby contaminated food samples were heat treated, inoculated with lactic acid bacteria and incubated. ‘Fermentation = No’ represents control groups (non-fermented foods), whereby contaminated food samples were heat treated and incubated with no inoculation. Results were collated across all contamination levels (0.5-50mg/kg), inoculation concentrations, incubation temperatures (23-45°C) and incubation times (5-72 hours). Error bars give 95% confidence interval.

The half-lives of organophosphate insecticides in apple juice, milk and wheat ranged from 9.5 hours to 21.0 days in fermented foods and ranged from 21.4 hours to 36.5 days in non-fermented foods. The shortest half-life of 9.5 hours was observed when pasteurised milk which had been spiked with dimethoate was inoculated with 10^9^ CFU/ml (10% v/v) *Lpb.plantarum* subsp*.plantarum* and incubated at 37°C for 24 hours (Yuan et al., 2021).

#### Species of Lactic Acid Bacteria

Considering all species of lactic acid bacteria and all insecticide types, fermentation with lactic acid bacteria led to a mean change in organophosphate insecticide half-life of -30.5%. The median percentage change in organophosphate insecticide half-life was negative for all species of lactic acid bacteria (Figure 5).

Single species of lactic acid bacteria that demonstrated strong potential for organophosphate insecticide degradation were *Lpb.plantarum* subsp*.plantarum, Lab.delbrueckii* subsp*.bulgaricus* and *Lvb. brevis;* where the median percentage change in organophosphate insecticide half-life was -42.3%, -25.0% and - 22.9%, respectively (Figure 4a), and the half-life in the presence of fermentation was shorter than the half-life in the absence of fermentation for all experiments. The combination of *S.thermophilus* and *Lab.delbrueckii* subsp*.bulgaricus* also demonstrated promising potential, with a median percentage change in organophosphate insecticide half-life of 45.9% (Figure 4a).

**Figure 4.**
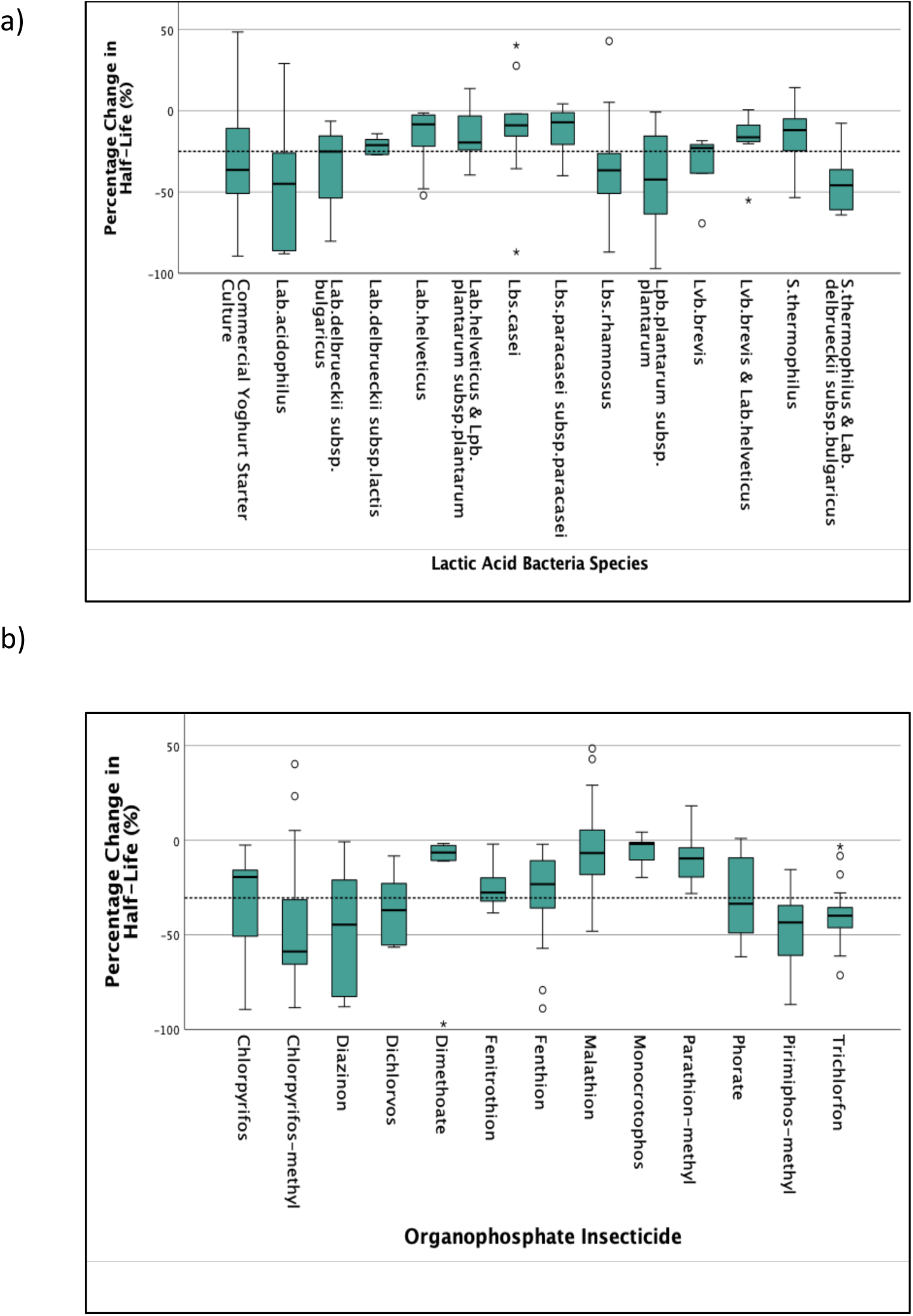
Percentage change in organophosphate half-life during controlled fermentation. Ten studies were analysed. The percentage change in half-life was based on the relative difference in half-life between treatment and control. The value is directional, with negative values representing a shortening of half-life. The unbroken horizontal lines within the boxes represent the medians for each insecticide, the box represents the interquartile range, the whiskers represent the minimum and maximum non-outlier values and the broken line across all values represents the mean for all insecticides. Results were collated across all food categories (apple juice, milk and wheat), all contamination levels (0.5-50mg/kg), inoculation concentrations, incubation temperatures (23-45°C) and incubation times (5-72 hours). Percentage change in organophosphate insecticide half-life was analysed by (a) Lactic acid bacteria species, independent of organophosphate insecticide type, and (b) Organophosphate insecticide type, independent of lactic acid bacteria species.

There were no lactic acid bacteria species that demonstrated poor insecticide degrading ability. Even for species such as *Lbs. casei* where the median percentage reduction in half-life was 9.0% (Figure 4a), fermentation with *Lbs. casei* could still shorten the half-life of diazinon in apple juice by up to 86.9% (Rezaei et al., 2021).

#### Organophosphate Insecticide Type

The median percentage change in organophosphate insecticide half-life was negative for all organophosphate insecticides (Figure 4b). Organophosphate insecticides that were observed to be particularly susceptible to enhanced degradation during fermentation were diazinon and pirimiphos-methyl; where the median percentage change in half-life was -44.6% and -43.4% (Figure 4b) and the half-life in the presence of fermentation was shorter than the half-life in the absence of fermentation for all experiments.

On the other hand, malathion was observed to be quite resistant to enhanced degradation during fermentation. The median percentage change in half life for malathion as a result of fermentation, was – 6.8% (Figure 4b), and in several experiments the half-life of malathion was extended as a result of fermentation (Bo et al., 2011; Zhang et al., 2014; Zhao et al., 2012; Zhou & Zhao, 2015). For example, Zhou and Zhao (2015) observed that when contaminated skim milk was inoculated with *Lab.acidophilus, Lbs.casei, Lbs.rhamnosus* or *S.thermophilus* the percentage change in half-life of malathion was 29.0%, 27.7%, 42.9% and 14.3%, respectively. In these experiments, fermentation extended the half-life of malathion.

### The Effect of Natural Fermentation on Organophosphate Insecticide Degradation is Uncertain

Three studies investigated organophosphate insecticide degradation during natural fermentation, and these were in cabbage and olives. In these studies, degradation did not always fit well with the first-order degradation model and therefore percentage degradation was used to compare organophosphate insecticide degradation rather than half-life.

#### Natural Fermentation

Two studies investigated organophosphate insecticide degradation during natural fermentation and compared this with no fermentation, showing mixed results (Kumral et al., 2020; Maden & Kumral, 2020).

During sauerkraut production, Maden and Kumral (2020) observed greater organophosphate insecticide degradation in vacuum-packed cabbage compared with fermented cabbage. After the 14-day fermentation, chlorpyrifos-methyl had degraded by 11.7% in the fermented cabbage and 68.9% in the vacuum-packed cabbage. Malathion had degraded by 58.6% in the fermented cabbage and 97.7% in the vacuum-packed cabbage. The greatest degradation for both organophosphate insecticides was observed in the vacuum-packed cabbage. Whilst the researchers measured lactic acid bacteria growth and pH in the fermented cabbage, this was not recorded for the vacuum-packed cabbage.

During olive production, Kumral and colleagues (2020)) observed greater organophosphate insecticide degradation in fermented olives compared with dehydrated black olives as well as natural black olives. By day 60, dimethoate had degraded by 66.3% in the fermented olives, 40.1% in dehydrated black olives and 10.3% in natural black olives. Whilst researchers did observe higher dimethoate degradation in the fermented olives, lactic acid bacteria were not detected in the fermented samples, and a pH decline was also not observed.

#### Natural Fermentation with Inoculation

Three studies investigated organophosphate insecticide degradation when natural fermentations were inoculated with *Lpb. plantarum* subsp*.plantarum* and compared this with no inoculation (i.e. natural fermentation only), showing mixed results. During pickled cabbage production, Zhou and colleagues (2015) observed greater organophosphate insecticide degradation in inoculated samples compared with uninoculated samples. After a 6-week fermentation, four organophosphates had degraded by 96.2% to 99.8% in inoculated samples and 80.6% to 93.1% in uninoculated samples. However, whilst a standard fermentation procedure was followed, there was no microbial analysis to confirm that lactic acid bacteria were present in the control samples, and pH was not measured.

A later study in cabbage observed greater degradation in inoculated samples for chlorpyrifos-methyl but not for malathion (Maden & Kumral, 2020). After a 14-day fermentation, chlorpyrifos-methyl had degraded by 30.7% in the inoculated samples and 11.7% in the uninoculated samples, whereas malathion had degraded by 34.0% in the inoculated samples and 58.6% in the uninoculated samples. Therefore, whilst inoculation with *Lpb. plantarum* subsp*.plantarum* increased the degradation of chlorpyrifos-methyl, it actually decreased the degradation of malathion.

In olives, inoculation did not influence dimethoate degradation (Kumral et al., 2020). After a 60-day fermentation, dimethoate had degraded by 66-68% in the inoculated samples and 66% in the uninoculated samples. Nevertheless, the inoculation did influence the microbial consortium, with lactic acid bacteria detected in the inoculated samples but not in the uninoculated samples.

### Organophosphate Insecticide Degradation during Fermentation with Lactic Acid Bacteria is affected by Contamination Level, Inoculum Concentration and Fermentation Temperature

Several factors influenced the level of organophosphate insecticide degradation that was achieved during fermentation (Table 2). Studies demonstrated that at the highest organophosphate insecticide contamination levels, there was reduced insecticide degradation during fermentation (Dordevic & Durovic-Pejcev, 2016; Rezaei et al., 2021; Yuan et al., 2021). Two studies observed that increasing the lactic acid bacteria inoculum concentration increased organophosphate insecticide degradation, with one of these studies observing that there may be an upper limit whereby further increases in inoculum results in no further degradation (Rezaei et al., 2021; Yuan et al., 2021). One study observed that different incubation temperatures led to significantly different levels of organophosphate insecticide degradation (Yuan et al., 2021).

**Table 2.**
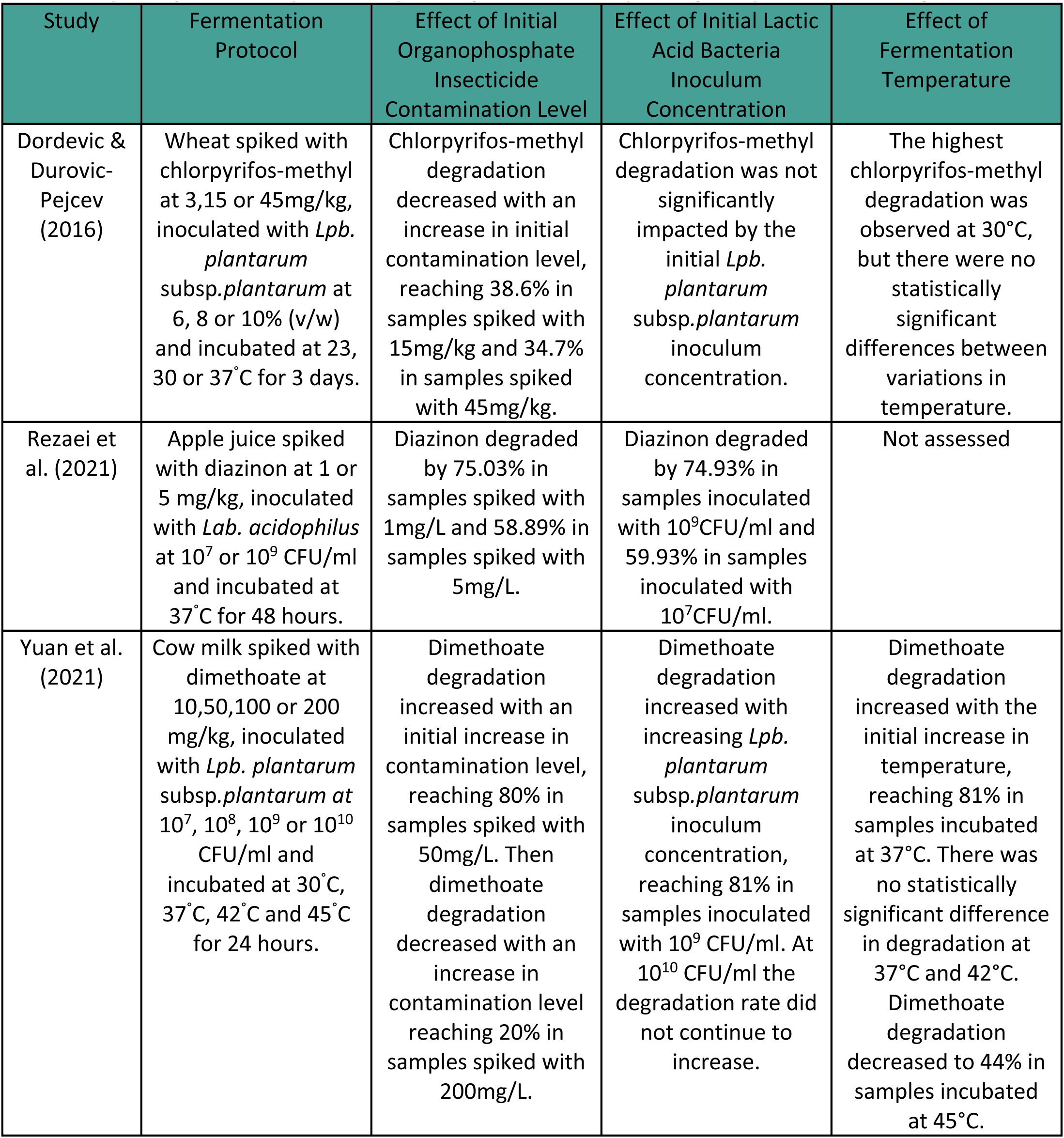
Parameters affecting organophosphate insecticide degradation during fermentation. Three studies investigated how different contamination levels and different fermentation parameters can affect organophosphate degradation. Initial organophosphate insecticide contamination level was the level of organophosphate insecticide spiked into the food in the lab which ranged from 1mg/kg to 200mg/kg. Initial lactic acid bacteria inoculum concentration was the level of lactic acid bacteria inoculated into the spiked food which was reported as either v/w or CFU/ml. Fermentation temperature was the temperature throughout the incubation period, which ranged from 23°C to 45°C. mg/kg = milligrams per kilogram. v/w = volume of lactic acid bacteria per weight of food, expressed as a percentage. CFU/ml = colony forming units per millilitre. °C = degrees Celsius.

| Study | Fermentation Protocol | Effect of Initial Organophosphate Insecticide Contamination Level | Effect of Initial Lactic Acid Bacteria Inoculum Concentration | Effect of Fermentation Temperature |
| --- | --- | --- | --- | --- |
| Dordevic & Durovic-Pejcev (2016) | Wheat spiked with chlorpyrifos-methyl at 3,15 or 45mg/kg, inoculated with *Lpb. plantarum* subsp.*plantarum* at 6, 8 or 10% (v/w) and incubated at 23, 30 or 37°C for 3 days. | Chlorpyrifos-methyl degradation decreased with an increase in initial contamination level, reaching 38.6% in samples spiked with 15mg/kg and 34.7% in samples spiked with 45mg/kg. | Chlorpyrifos-methyl degradation was not significantly impacted by the initial *Lpb. plantarum* subsp.*plantarum* inoculum concentration. | The highest chlorpyrifos-methyl degradation was observed at 30°C, but there were no statistically significant differences between variations in temperature. |
| Rezaei et al. (2021) | Apple juice spiked with diazinon at 1 or 5 mg/kg, inoculated with *Lab. acidophilus* at 10<sup>7</sup> or 10<sup>9</sup> CFU/ml and incubated at 37°C for 48 hours. | Diazinon degraded by 75.03% in samples spiked with 1mg/L and 58.89% in samples spiked with 5mg/L. | Diazinon degraded by 74.93% in samples inoculated with 10<sup>9</sup>CFU/ml and 59.93% in samples inoculated with 10<sup>7</sup>CFU/ml. | Not assessed |
| Yuan et al. (2021) | Cow milk spiked with dimethoate at 10,50,100 or 200 mg/kg, inoculated with *Lpb. plantarum* subsp.*plantarum* at 10<sup>7</sup>, 10<sup>8</sup>, 10<sup>9</sup> or 10<sup>10</sup> CFU/ml and incubated at 30°C, 37°C, 42°C and 45°C for 24 hours. | Dimethoate degradation increased with an initial increase in contamination level, reaching 80% in samples spiked with 50mg/L. Then dimethoate degradation decreased with an increase in contamination level reaching 20% in samples spiked with 200mg/L. | Dimethoate degradation increased with increasing *Lpb. plantarum* subsp.*plantarum* inoculum concentration, reaching 81% in samples inoculated with 10<sup>9</sup> CFU/ml. At 10<sup>10</sup> CFU/ml the degradation rate did not continue to increase. | Dimethoate degradation increased with the initial increase in temperature, reaching 81% in samples incubated at 37°C. There was no statistically significant difference in degradation at 37°C and 42°C. Dimethoate degradation decreased to 44% in samples incubated at 45°C. |

## Discussion

This review aimed to bring together the evidence on the effectiveness of lactic acid fermentation in reducing organophosphate insecticide residues in food and explore the factors that affect degradation. Data on organophosphate insecticide degradation during lactic acid fermentation was analysed and organophosphate insecticide half-lives in fermented and non-fermented foods were estimated. Results demonstrated that, on average, all organophosphate insecticides degraded over time, irrespective of fermentation. This finding demonstrates how important it is that studies in this area have a negative control. Negative controls help to establish whether insecticide degradation is a result of fermentation or due to inherent degradation in the food matrix over time. Only studies that used a negative control were included in our analysis and we were able to establish that, in most cases, controlled fermentation could speed up the degradation of organophosphate insecticides in food, beyond the rate of degradation in the food matrix, reflected in shorter half-lives. Whilst in some cases this trend was also observed in natural fermentations, there were fewer studies and less consistent results.

### Quality of included studies

Included studies were assessed for Risk of Bias (RoB) using the RoBDMAT tool which was originally developed for studies in dentistry (Delgado et al., 2022). Whilst the domains and signalling questions in this tool could be adapted to studies in food science and microbiology, it is recommended that in the future a tool is developed specifically for these types of laboratory-based studies.

The RoB assessments of included studies revealed that there were sources of potential bias. All studies were unblinded and several studies underreported the sample size rationale and randomisation process. Several studies did not adequately report on statistical analysis which may have been a reflection of differences in study objectives and hypotheses. It is recommended that future studies are blinded, provide a sample size rationale, provide sufficient detail on the randomisation process and conduct statistical analysis to validate observed differences between groups.

### Mechanisms of Organophosphate Insecticide Degradation during Fermentation

There are several mechanisms which may account for the increased degradation rate of organophosphate insecticides during fermentation. Insecticides degrade through non-biological (abiotic) and biological (biotic) process. Whilst abiotic degradation refers to the chemical and mechanical breakdown of insecticides through reactions such as hydrolysis, biotic degradation refers to the transformation of the insecticides by microorganisms (Kiruthika et al., 2025; Sarlak et al., 2021).

#### Abiotic Degradation

Organophosphate insecticides are susceptible to hydrolysis, with the rate of hydrolysis being influenced by several factors, including pH. This concept is important in fermentation studies because, during the process of lactic acid fermentation, the pH of food becomes more acidic as lactic acid is produced (Wang et al., 2021). Less than half of the studies in this review measured pH, and only two studies measured the pH in both treatment and control samples (Maden & Kumral, 2020; Rezaei et al., 2021). Nonetheless, studies demonstrated a decline in pH over time with the pH dropping as low as 3.9 in fermented apple juice (Rezaei et al., 2021), 3.2 in fermented milk (Yang et al., 2024), 3.4 in fermented cabbage (Maden & Kumral, 2020) and 4.0 in fermented olives (Kumral et al., 2020). It is well established that the majority of organophosphate insecticides, aside from diazinon, degrade rapidly in alkaline conditions (Turner, 2024). In the pH range of 5-7 many organophosphate insecticides are relatively stable, but less is known about what happens at the lower pH values observed during lactic acid fermentation (Turner, 2024). In order to understand the contribution of hydrolysis to organophosphate insecticide degradation during fermentation, further research should ensure that pH is measured throughout the fermentation process, and that there is consideration for pH adjusted controls.

#### Biotic Degradation

Several microorganisms are able to degrade organophosphate insecticides through enzyme-mediated processes (Armenova et al., 2023). Therefore, organophosphate insecticide degradation during fermentation with lactic acid bacteria may be augmented by enzymatic breakdown. This would involve the genes encoding for organophosphate degrading enzymes being present and active in the lactic acid bacteria during fermentation. Our analysis demonstrated that the rate of organophosphate insecticide degradation may be influenced by the species of lactic acid bacteria present in the fermentation, with several lactic acid bacteria demonstrating promising potential. *Lab.delbrueckii* subsp*.bulgaricus* is one such species which has been the subject of further analysis. Yang and colleagues (2024) analysed the genes and enzymes responsible for organophosphate insecticide degradation in their study involving chlorpyrifos-spiked milk and *Lab. delbrueckii* subsp*.bulgaricus*. Through RNA-seq transcriptomic analysis, the researchers analysed changes in gene expression during fermentation. Of the 269 upregulated genes, several related to the degradation of chlorpyrifos, including five belonging to the phosphoesterase / diphophatase class of genes and ten belonging to the hydrolase class of genes. More research is needed to better understand whether the genes that encode organophosphate insecticide degrading enzymes are present and active in the common lactic acid bacteria present in food fermentations.

### Organophosphate Insecticide Degradation Products

The goal of organophosphate insecticide degradation is to yield compounds which are non-toxic or less toxic than the original compound and have less of an impact on human health and the environment (Leskovac & Petrović, 2023). However, depending on the specific degradation pathway, there is potential for organophosphate insecticides to form intermediate degradation products that are more toxic than the parent compound and persist in the environment (Jaiswal et al., 2024; Kiruthika et al., 2025; Sarlak et al., 2021). For example, in insects and mammals, oxidation of the P-S bond in malathion produces malaoxon which is several times more toxic than malathion (Jaiswal et al., 2024). Despite this knowledge, few studies in food fermentations measure degradation products. Of the 14 studies included in this review, only two measured degradation products (Yang et al., 2024; Yuan et al., 2021). During milk fermentation, dimethoate transformed into several intermediate degradation products including omethoate, whereas chlorpyrifos yielded products such as 3,5,6-trichloro-2-pyridinol (TCP). Omethoate is known to be several times more toxic to humans that dimethoate (Office of Chemical Safety and Environmental Health, 2007). TCP is also toxic to humans and is more persistent in the environment than its parent compound (Rivero et al., 2022). It is important that future studies in this area measure organophosphate degradation products, to firstly gain a better understanding of degradation pathways and secondly to ensure that degradation results in less toxic compounds.

### Susceptibility of Organophosphate Insecticides to Enhanced Degradation

Whilst some organophosphate insecticides were observed to be quite susceptible to enhanced degradation during fermentation, others were more resistant. This may be attributed to varying physical and chemical characteristics (Sarlak et al., 2021). In particular, the nature of the atoms that are attached to the central phosphorus atom (commonly oxygen or sulphur) as well as the structure of the leaving and non-leaving groups, may be contributing factors (Jaiswal et al., 2024; Sarlak et al., 2021; Silva & Orth, 2023). Our analysis indicated that malathion may be resistant to enhanced degradation during fermentation. Malathion is a phosphorodithioate (containing a P=S bond) with an aliphatic leaving group, and these factors may have contributed to slower reactivity (Silva & Orth, 2023; Turner, 2024). A study in MRS medium also found that malathion was resistant to enhanced degradation following inoculation with lactic acid bacteria. Nonetheless, studies in soil bioremediation have found that microbial consortiums including *Micrococcus aloeverae, Bacillus cereus,* and *Bacillus paramycoides* have successfully degraded malathion (Kosimov et al., 2025). Further research is needed to better understand the structure-reactivity relationship of organophosphate insecticides during fermentation.

### Degradation Kinetics

In our analysis, we calculated degradation rate constants, and subsequently half-lives of organophosphate insecticides, based on first-order degradation kinetics. However, there were indicators that insecticide degradation during fermentation diverged from first-order kinetics in some experiments. Several studies demonstrated that the initial organophosphate insecticide contamination level impacted the level of degradation during fermentation, which is not consistent with a first-order degradation model. A general trend was observed that at the highest contamination levels there was lower pesticide degradation (Leskovac & Petrović, 2023). This may have related to the saturation of enzyme active sites as insecticide concentration increased (Wirsching et al., 2020). This may also have related to the inhibition of lactic acid bacteria at high insecticide concentrations. This is supported by evidence from Li and colleagues (2018) who monitored the growth of *Lpb.plantarum* subsp*.plantarum* in culture with and without phorate. Whilst the researchers observed phorate degradation, they also observed that the bacteria grew significantly slower after 12 hours in the presence of phorate. *Lpb.plantarum* subsp*.plantarum* in culture containing the highest concentrations of phorate also showed significantly lower viable counts. We also observed that in natural fermentations, organophosphate insecticide degradation did not always follow the pattern of exponential decay which defines the first-order model (Kumral et al., 2020; Maden & Kumral, 2020). Whilst most studies in this area assume pseudo-first order degradation kinetics, it is important that further research is conducted to better understand the true degradation behaviour of organophosphate insecticides during food fermentations, at both high and low contamination levels, and in controlled and natural fermentations, so that the most reliable models are used.

## Conclusion

In most cases, controlled fermentation with lactic acid bacteria shortened the half-life of organophosphate insecticides in food. Degradation may have been a result of abiotic or biotic mechanisms, or their combined influence. The species of lactic acid bacteria involved in fermentation may impact the rate of insecticide degradation, with *Lpb.plantarum* subsp*.plantarum, Lab.delbrueckii* subsp*.bulgaricus* and *Lvb. brevis* showing promising potential. The rate of degradation may relate to the type of organophosphate insecticide under investigation, with insecticides such as malathion being potentially more resistant to degradation. It may also relate to the initial contamination level, whereby high contamination levels may inhibit the growth of lactic acid bacteria and subsequently reduce their insecticide-degrading ability.

The collated data demonstrated that controlled fermentation with lactic acid bacteria is an effective method to reduce organophosphate insecticide residues in food. This review has increased our understanding of fermentation as a simple, safe, low-cost option for reducing organophosphate insecticides in the food supply. This new knowledge may assist food manufacturers in creating products with reduced organophosphate insecticide residues and help to grow the market for fermented foods. For consumers who are trying to reduce their exposure to pesticides, this knowledge may help them to make more informed food choices. Having a better understanding of the factors that can impact organophosphate insecticide degradation during fermentation may also lead to the development of more accurate processing factors for fermentation, informing more reliable pesticide exposure assessments, and protecting public health.

