## Supplementary Figure S1 for "Analysis of Organophosphate Insecticide Half-Lives in Foods Fermented with Lactic Acid Bacteria"

### Supplementary Information

(a)

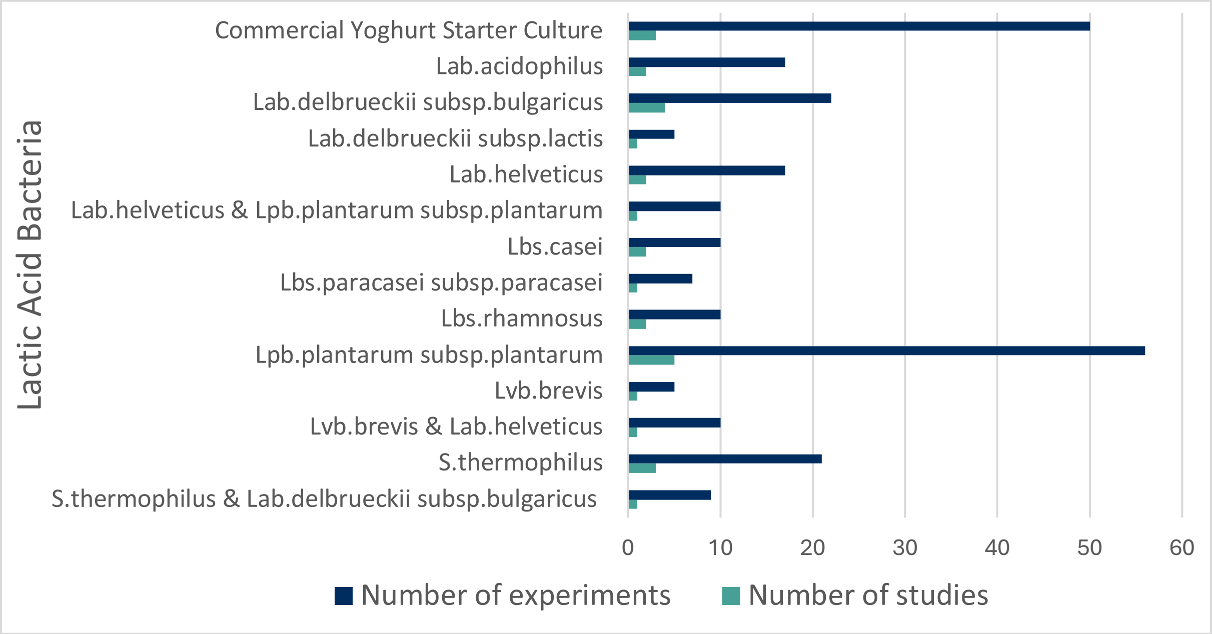


(b)

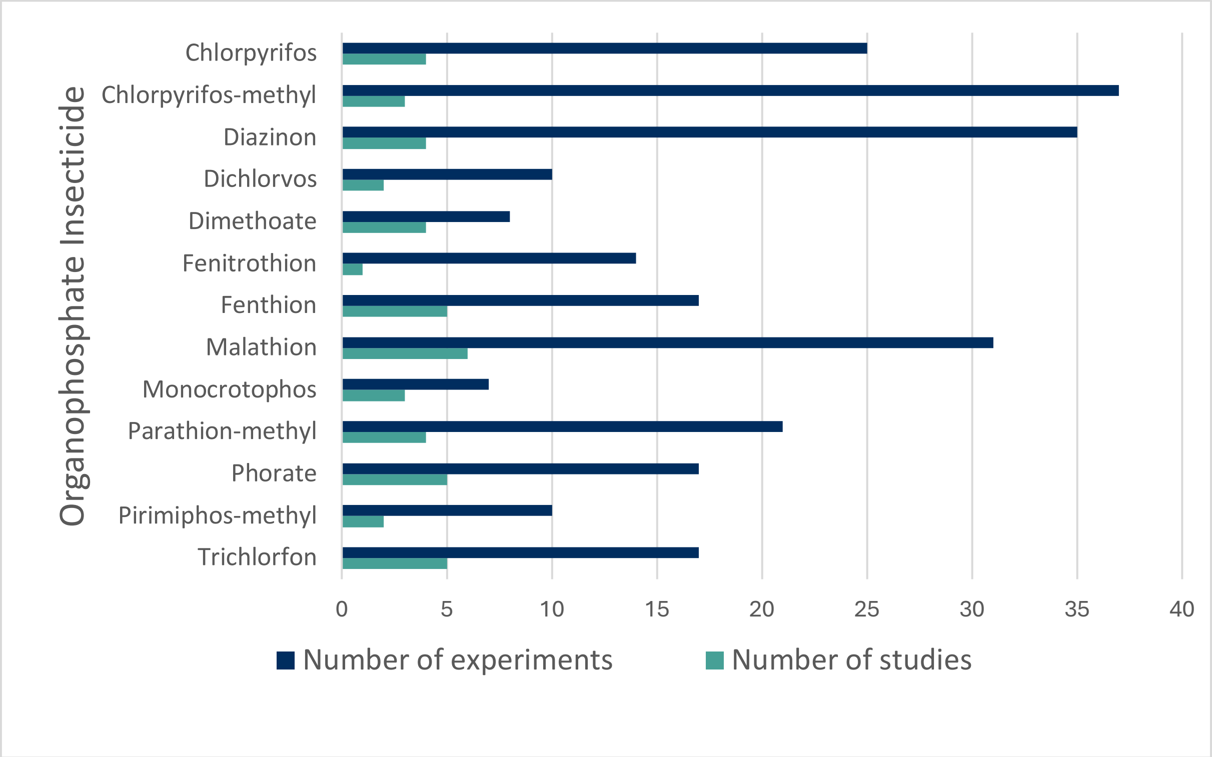


**Figure S1. Number of studies and treatments included in the analysis for controlled fermentations.** 13 studies involving 249 treatments were included in the analysis. A controlled fermentation was described as a fermentation process involving initial heat treatment followed by deliberate inoculation with lactic acid bacteria and subsequent incubation. These studies were carried out in apple juice, milk and wheat. The number of studies and treatments for each lactic acid bacteria, and combination of lactic bacteria, are shown in (a) and for each organophosphate insecticide are shown in (b).
